# Effects on cell viability, growth and morphology of *C. albicans* SC5314 biofilms after kINPen®09 plasma treatment

**DOI:** 10.1101/323600

**Authors:** O. Handorf, T. Weihe, S. Bekeschus, A. C. Graf, U. Schnabel, K. Riedel, J. Ehlbeck

**Author notes:** Corresponding author: Oliver Handorf, <u>Leibniz Institute for Plasma Science and Technology, 9 (INP)</u> Felix-Hausdorff-Str. 2, 17489 Greifswald, Germany.

## Abstract

Microorganisms are predominantly organized in biofilms, where cells live in dense communities and are more resistant to external stresses compared to their planktonic counterparts. With *in vitro* experiments, the susceptibility of *Candida albicans* biofilms to a non-thermal plasma treatment (plasma source kINP®09), in terms of growth, survival, and cell viability was investigated. Behind that background, the *C. albicans* strain SC5314 (ATCC® MYA-2876™) was plasma treated for different time periods (30 s, 60 s, 120 s, 180 s, 300 s). The results of experiments embracing colony forming units, fluorescence LIVE/DEAD assays, and XTT assays revealed a negative influence of the plasma treatment on the proliferation ability, vitality, and the metabolism of *C. albicans* biofilms, respectively. Morphological analysis of plasma-treated biofilms using atomic force microscopy supported the indications for lethal plasma effects concomitant with membrane disruptions and the loss of intracellular fluid. Controversial to other publications, fluorescence- and confocal laser scanning-microscopic inspection of plasma-treated biofilms indicated, that an inactivation of cells mainly appeared on the bottom side of the biofilms. If this inactivation leads to a detachment of the biofilms from the overgrown surface, it might offer completely new approaches in the plasma treatment of biofilms. Because of its biochemical-mechanical mode of action, resistances of microbial cells against plasma are unknown at this state of research.

## Importance

Microbial communities are an increasing problem in medicine but also in industry. Thus, an efficient and rapid removal of biofilms is becoming increasingly important. With the aid of the kINPen®09, decisive new findings on the effects of plasma on *C. albicans* biofilms could be obtained. This work was able to show for the first time that the inactivation of biofilms mainly takes place on the bottom side, which in turn offers new possibilities for the removal of biofilms by other strategies, e.g. mechanical treatment. This results demonstrated that non-thermal atmospheric pressure plasma is well-suited for biofilm decontamination.

## Introduction

*Candida albicans* is one of the most common pathogens causing mycosis worldwide. It is the eponym of the *Candida* group, which is a yeast genus that can cause fungal infections in humans and animals. *C. albicans* is a member of the *Fungi imperfecti*, that multiplies by budding (1). Typically, *C. albicans* cells bind to biogenic or abiogenic surfaces and grow as three-dimensional structures also designated as biofilms (2). The fungus can form mono- or multi-species biofilms which can be considered as densely packed cellular communities embedded in an extracellular matrix (3). These biofilm communities are known to be more resistant to physical and chemical stresses compared to planktonic cells, and can cause major economic loss within industrial sectors (4). Moreover, more than four million people in Germany suffer from chronic wounds, which are often caused by resistant biofilm-forming pathogens (5). The treatment costs of these patients amount up to four billion euro per year only in Germany (6). Catheter-related bloodstream infections (CRBSI), caused by biofilms and there direct access to the bloodstream, are a big problem in the United States. Several studies counted 80.000 CRBSI each year in the United States with an estimation of 4.888$ to 11.591$ for each CRBSI event (7, 8). Another important factor is the biofilm-associated infection of implants such as prosthetic hips and knees. A study revealed the demand for primary total hip arthroplasties is estimated to grow by 174 % to 572.000 and the demand for primary total knee arthroplasties is projected to grow by 673 % to 3.48 million procedures between 2005 and 2030 in the United States (9). These facts demonstrate the importance of the successful treatment of biofilms in medicine.

*Candida* sp. is responsible for two types of infections: superficial infections like oral and vaginal candidiasis, and systemic infections (10, 11). In about 75 % of the world’s population, *C. albicans* appears in the oral cavity and is the fourth most common cause of hospital-acquired systemic infections in the United States with up to 50 % mortality (12–15). Due to the increased lifespan of today’s population, there is also an upraise in the number of patients who wear dental prostheses, which leads to a concomitant increase in *Candida* stomatitis (16–18). Dental diseases like periodontitis, gingivitis, carious, oral candidiasis, or periimplantitis are relevant diseases for people of all ages. Depending on the oral food intake, the oral cavity is constantly exposed to a high level of microbial contamination induced by the assimilated food (19). Thus, the germ load of the oral cavity is much higher than for other parts of the human body. Lesions in the oral cavity often lead to diseases such as oral submucous fibrosis or stomatitis (20) and therefore, play an important role as a symptom for immunodeficiency. During the infection process, colonization factors such as adhesins, invasins, hyphae and its thigmotropic properties are of major importance (21). Together with *Staphylococcus aureus, C. albicans* has a much higher incidence of 11-65 % for a stomatitis than in a mono-species biofilm (22). At present, the conventional treatment method of biofilms are antibiotics. Biofilms have a higher resistance to antimicrobial agents and a higher physico-mechanical tolerance (23, 24). These resistances are mainly due to complex interactions of the cells in the biofilm and the surrounding extracellular matrix (3, 25–27).

In 243 episodes of candidaemia, 45 (19 %) had reduced susceptibility (and 27 of them were fully resistant) to fluconazole, the main antifungal agent for candidaemia treatment. Despite 8 % of *C. albicans*, 4 % of *C. tropicalis* and 4 % of *C. parapsilosis* had reduced susceptibility to fluconazole, these species embrace 36 % of the reduced-suspectibility group and 48 % of the fully resistant group (28). Thus, there is a much higher probability for fully resistant candida albicans strains than for strains with reduced susceptibility. Resistance of *Candida* sp. to azole antifungal agents can occur via 4 different signaling pathways. On the one hand by increased release of azoles by upregulation of the multidrug-efflux transporter genes or by amino acid substitution in Erg11p. On the other hand by upregulation of ERG11 or an alteration of the ergosterol biosynthetic pathway (29). Antimicrobial resistance will be of increasing interest to society, as it is associated with rising mortality rates and higher medical costs (30). Therefore, alternatives to conventional antibiotic therapies are urgently needed.

A new innovative technology addressing these problems is non-thermal atmospheric pressure plasma (NTP). With moderate temperatures, NTPs avoid thermal host-tissue damage, but have in fact much higher antimicrobial effects than e.g. UV- or hydrogen peroxide-treatments (31, 32). In addition, NTPs are very effective against antibiotic-resistant pathogens and there are no antimicrobial resistance mechanisms against plasma known to date (33–35). Therefore, NTPs could be used for the decontamination of dental cavities and a wide range of other applications and devices in industry and medicine (36).

A basic understanding of the mechanistic effects of plasma on pro- and eukaryotic biofilms is a prerequisite for its application. Consequently, this work investigated the inactivation effects of plasma on *C. albicans* biofilms. For this purpose, biofilm viability, vitality and metabolism were examined after plasma treatment. The impact of plasma treatment on the three-dimensional biofilm structures was examined employing epifluorescence and confocal laser scanning microscopy (CLSM), moreover, cell morphological changes were detected with atomic force microscopy (AFM).

## Results

The antimicrobial effect of plasma components on biofilm cells might be caused by different mechanisms. To unravel the exact mode-of-action of the plasma treatment, it’s effect on *C. albicans* biofilms was monitored using colony forming unit (CFU)-counting to investigate the proliferation of the surviving cells (37), vitality assays with fluorescence LIVE/DEAD™ staining and a metabolism assay using XTT.

**Effects of plasma treatment on the proliferation ability of biofilm cells**. Five different treatment times in between 30 and 300 s were selected, which should demonstrate the dynamic range of the plasma effect. Six biofilms were treated with plasma for the respective time in fourfold repetition **(Fig. 1)**. The effects of plasma treatment on the biofilm CFU is displayed as reduction factor (RF). After 60 s plasma treatment time, a significant increase in the RF has been detected. The RF reached a maximum of 1.83 after 180 s plasma treatment. Surprisingly, longer treatment times led to the same or a slightly decreased RF.

**Figure 1:**
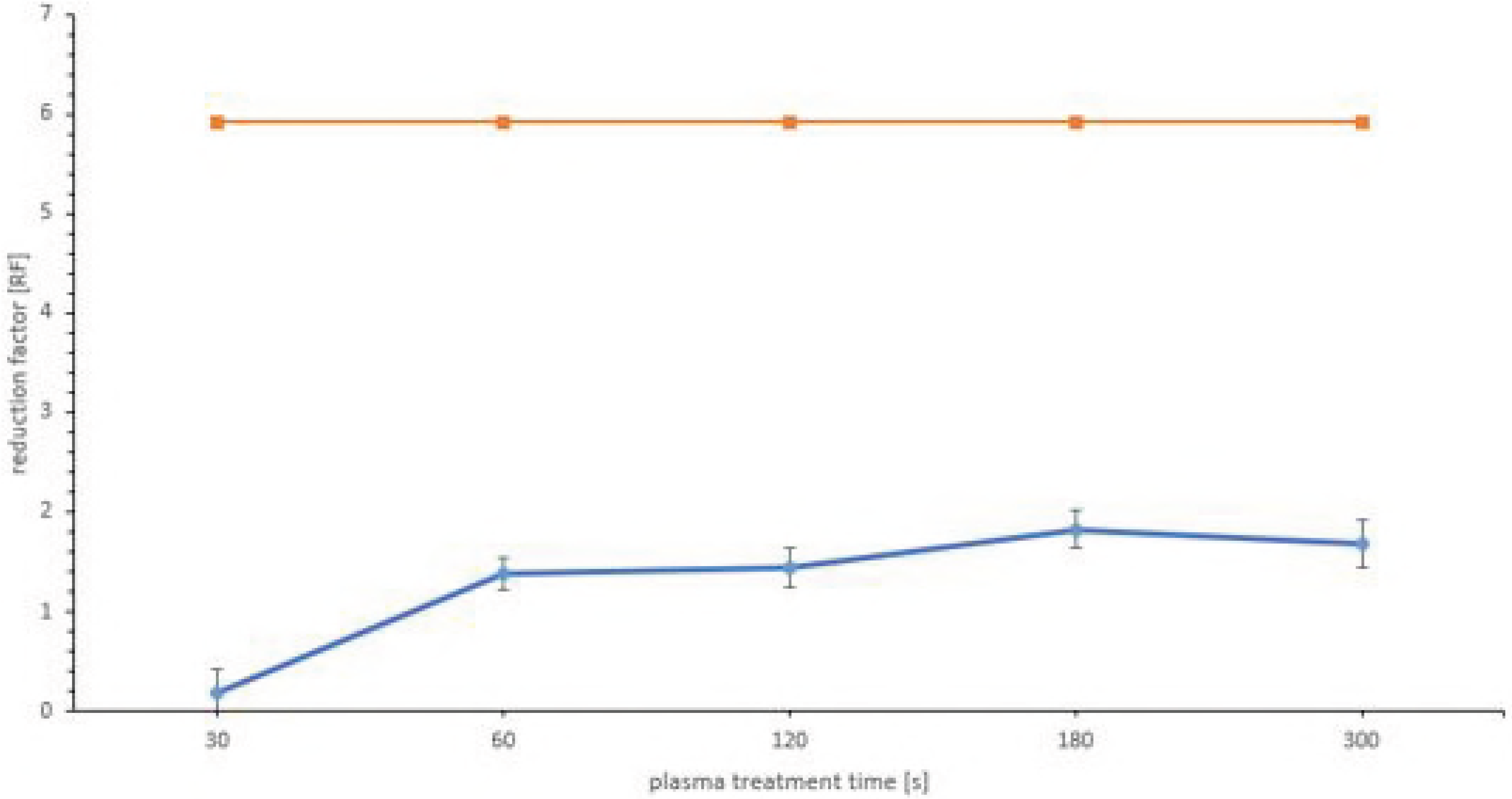
Detection limit (orange line). *Candida albicans* SC5314 reduction factor (RF) after 30, 60, 120, 180 and 300 s plasma treatment time with the kINPen™09 (blue line). The CFU was calculated using the formula 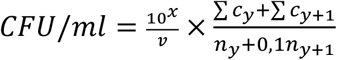. The RF was calculated using the formula *RF = MV_K_log_10_ − MV_P_log_10_* The data points are the mean value of the total population of the treatment time from the quadruple repetition. The error bars of the data points are the confidence reference (CR) of the treatment time calcualated with the formula 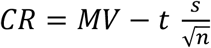 No significance was given in the results intersecting the x-axis (30 s of treatment). p < 0.05.

**Effects of plasma treatment on the biofilm vitality**. The LIVE/DEAD™ BacLight™ assay kit (Thermo Scientific, Waltham, USA) contains the DNA intercalating dyes SYTO™9 staining living cells of the biofilm and propidium iodide, which is membrane impermeable and reduces SYTO™9 in cells with damaged cell membranes. After 30 s of plasma treatment, a decrease in the ratio G/R of 0.97 was detected, which corresponded to a 39 % reduction **(Fig. 2)**. After 60 s of plasma treatment, a maximum decrease of 1.78 in the ratio G/R was observed, which corresponded to a 71.08 % reduction. There was no significant change in the ratio G/R after longer treatment times **(Fig. 2)**. Thus, significant changes in the vitality of the biofilm took place only in the first 60 s of plasma treatment.

**Figure 2:**
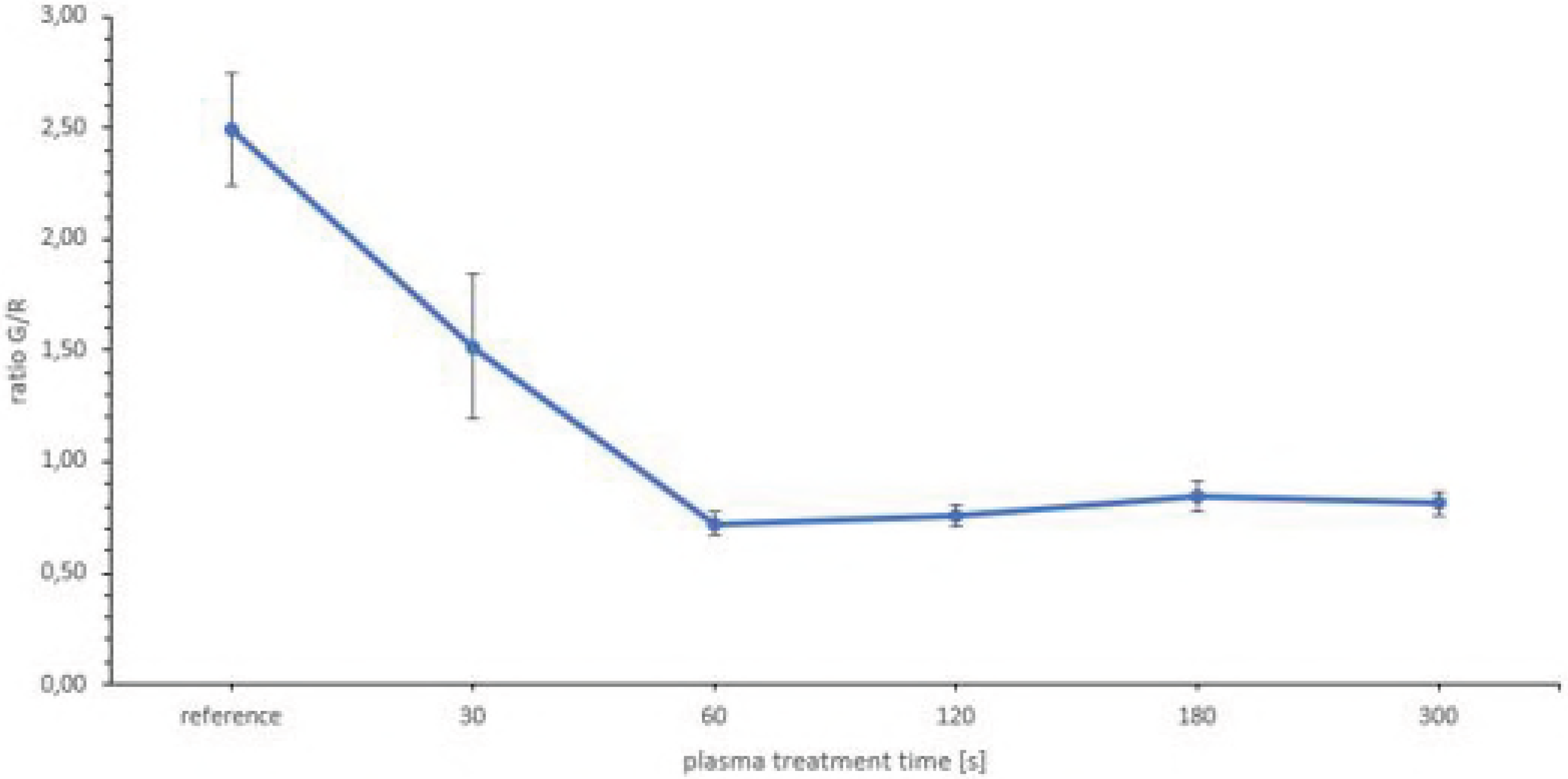
Fluorescence measurement of *Candida albicans* SC5314 biofilms after 30, 60, 120, 180 and 300 s plasma treatment with the kINPen™09. The ratio G/R is defined as the division of the extinction of green fluorescence by the extinction of red fluorescence. The data points are the mean value of the total population of the treatment time from the quadruple repetition. The error bars of the data points are the CR of the treatment time calcualated with the formula 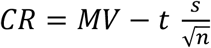 p < 0.05.

**Effect of plasma treatment on the metabolism of the cells.** Due to the fact, that the CFU and the LIVE/DEAD™-fluorescence showed antimicrobial effects after plasma treatment, the metabolism of cells was investigated as a third indicator of inactivation. For this, the XTT assay was used. Until 60 s of plasma treatment, a continuous decrease in the metabolism of the cells was observed **(Fig. 3)**. After 60 s, the extinction was reduced by 1.21. Because the extinction of the XTT dye is corresponding to the metabolic activity of the cells, this value equaled a reduction of 89 % in the cell metabolism. Longer treatment times lead to no significant alterations in cell metabolism compared to the 60 s plasma treatment time **(Fig. 3)**.

**Figure 3:**
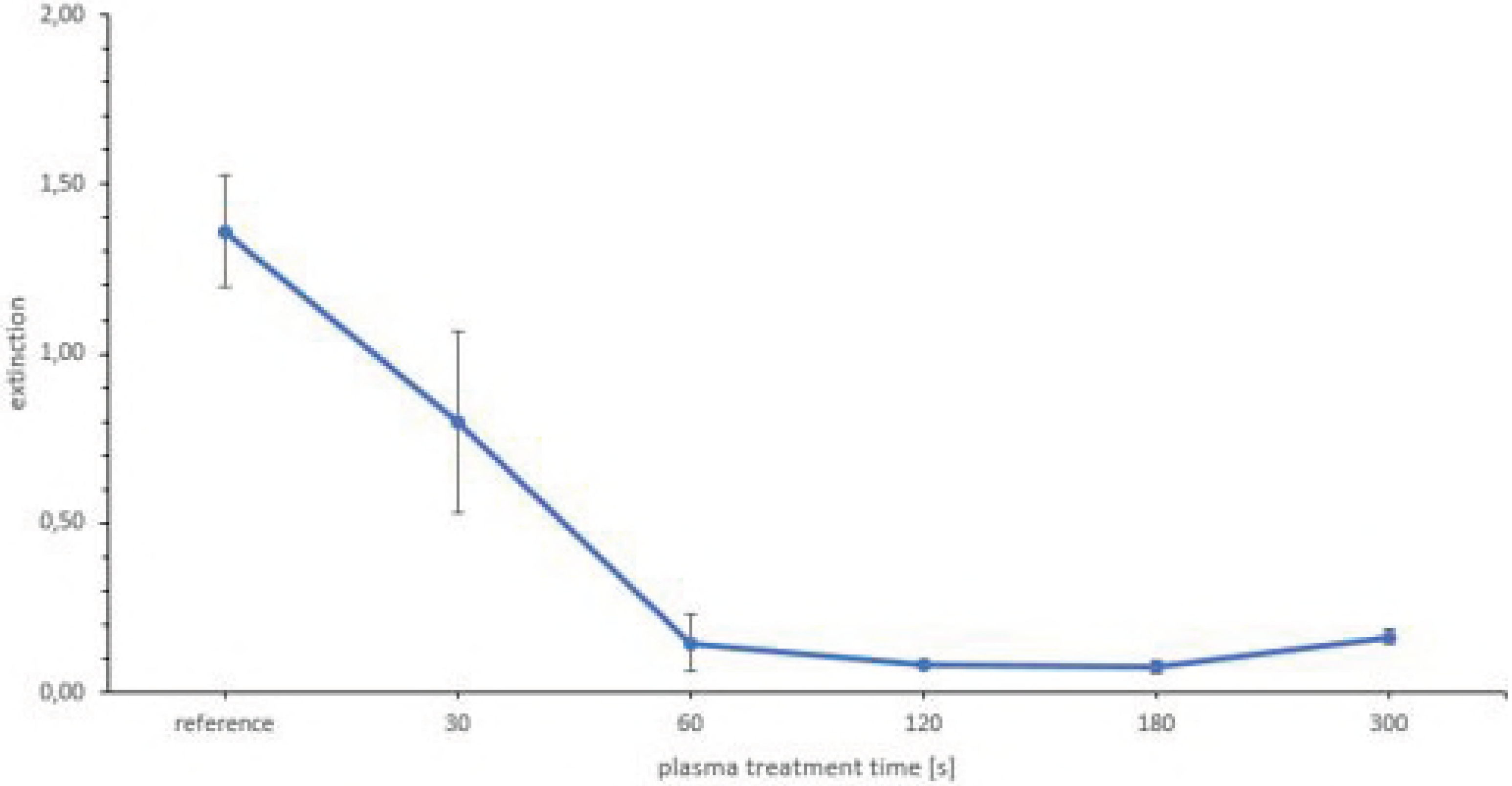
XTT assay as a metabolic measurement of *Candida albicans* SC5314 biofilms after 30, 60, 120, 180 and 300 s plasma treatment with the kINPen™09. The data points are the mean value of the total population of the treatment time from the quadruple repetition. The error bars of the data points are the CR of the treatment time calcualated with the formula 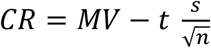. p < 0.05.

**Confirmation of plasma treatment effects by fluorescence microscopy.** The LIVE/DEAD™ BacLight™ staining was also used for an epifluorescent inspection of plasma-treated *C. albicans* biofilms cultivated in 96-well plates. Bottom view images of the biofilms revealed, that 30 s of treatment mainly affects the area of the biofilm that was in the direct range of the plasma effluent **(Fig. 4B)**. This area showed orange/yellow staining representing dead cells, which was not observed in the untreated control biofilms. With increasing treatment time, the effect could be observed throughout the entire well **(Fig. 4C-F)**. After 120 s treatment time, living cells were visible in the center of the biofilm **(Fig. 4D)**. This effect increased with longer treatment times.

**Figure 4:**
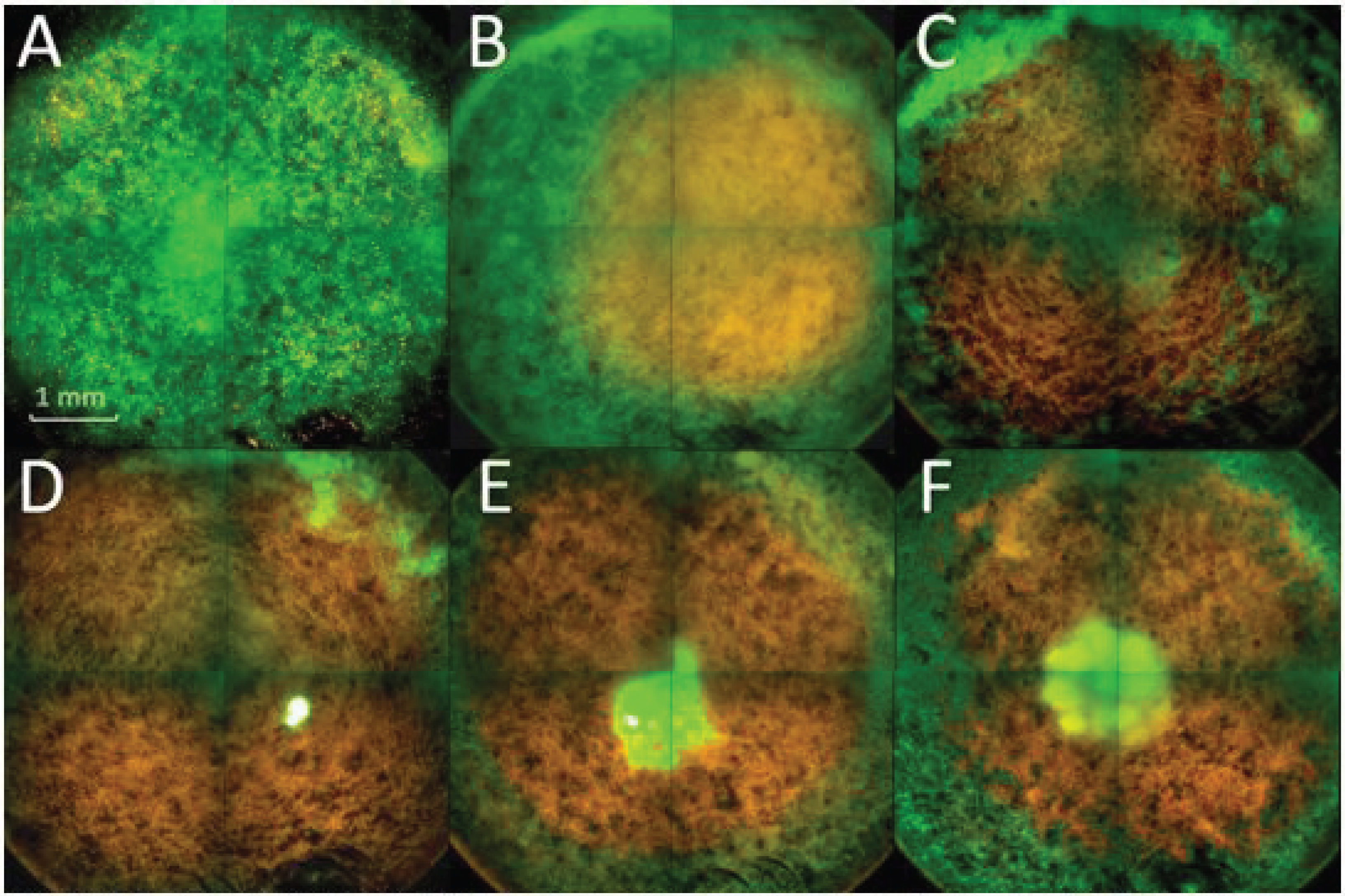
Overview with the Operetta CLS™ fluorescence microscope of plasma-treated biofilms. A) reference; B) 30 s plasma treatment time; C) 60 s plasma treatment time; D) 120 s plasma treatment time; E) 180 s plasma treatment time; F) 300 s plasma treatment time. All samples were treated with the kINPen™09. The biofilms were stained with SYTO™9 as green fluorescence (living) and propidium iodide as red fluorescence (dead).

**CLSM confirmed cell inactivation being most dominant on the bottom of the biofilms.** Using CLSM, 5.5 μm sections of the treated biofilms were acquired and displayed in orthogonal and 3D view **(Fig.5)**. All plasma treatments caused an enhanced propidium iodide staining indicating cell death compared to the reference **(Fig. 5**, **orthogonal view)**. After 30 s, plasma treatment effected mostly cells located at the bottom of the biofilm, indicated by a strong propidium iodide staining, whereas the top of the biofilm appeared to be less affected **(Fig. 5**, **3D-view)**.

**Figure 5:**
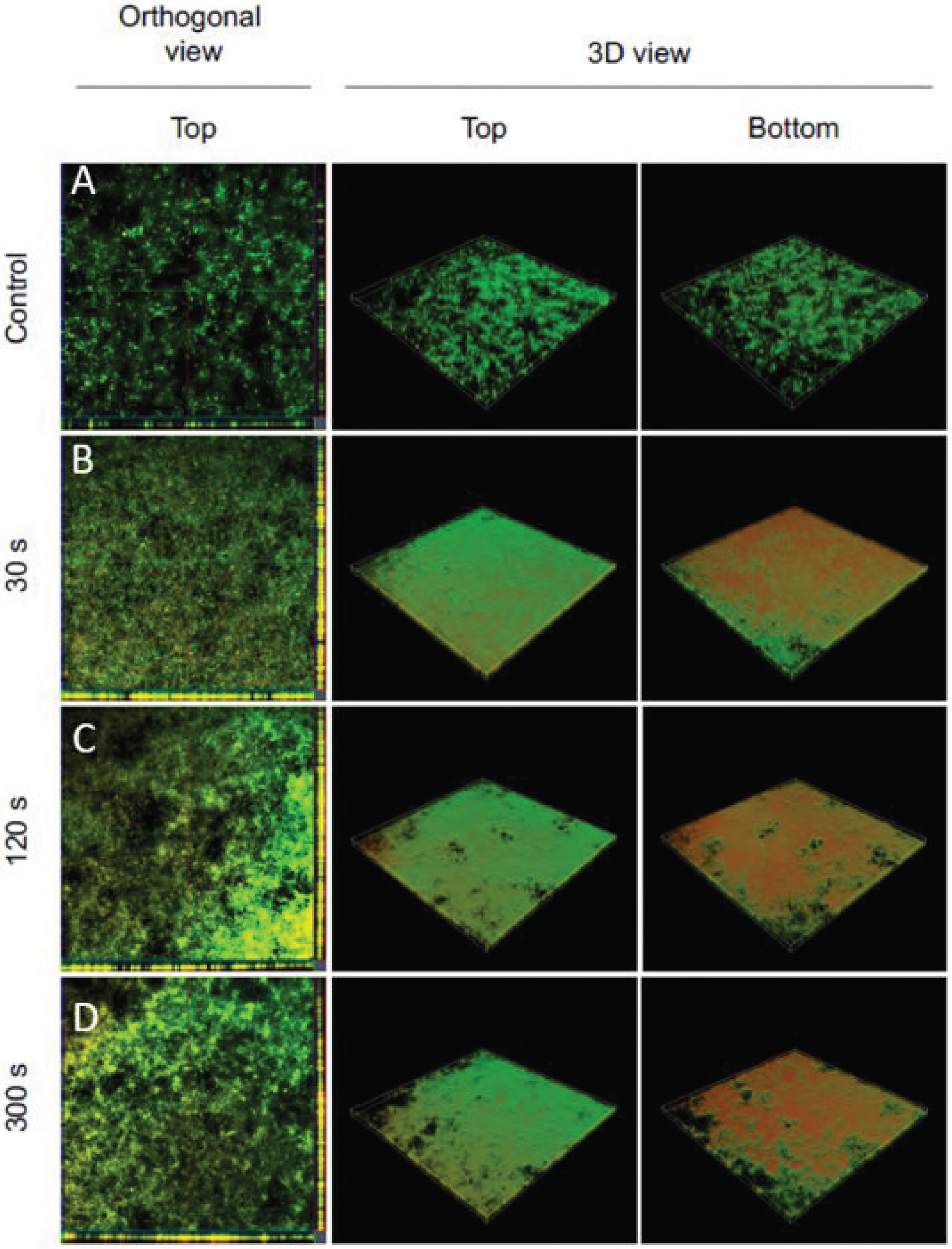
confocal-laser-scanning-mircroscopy (CLSM) of plasma-treated biofilms with the kINPen™09. A) reference B) 30 s plasma treatment time; C) 120 s plasma treatment time; D) 300 s plasma treatment time. All images were taken with the Zeiss LSM 510™ using an argon laser and a 10x objective (air NA = 0.3). The following measurement settings were used: Average = 4; Speed = 7; Pinhole (red) = 52.0; Pinhole (green) = 70.0; Digital Offset = -0.10; Gain (red) = 850; Gain (green) = 650 and a laser intensity of 10 % for both colors. The figure shows 3D images of the biofilms, but the images are rotated horizontally by 180°. The left side of the image shows an orthogonal view of the top layer of the biofilm. The center of the image represented the top of the biofilm and the right side of the image represented the bottom of the biofilm.

**AFM confirmed alterations in the cell morphology of the biofilm after plasma treatment.** AFM is often used to investigate cell morphological alterations in the μm range. **(Fig. 6)** showed AFM images of an untreated biofilm and plasma-treated biofilms for different times periods. Each image was taken as a topographic image on the left and as an error signal image on the right. In order to clearly detect the alterations caused by the plasma treatment, the average treatment time and the maximum treatment time were examined and recorded. **(Fig. 6)** indicates major changes in the cell morphology even after 120 s treatment time. Membrane disruptions and the loss of intracellular fluid could be detected. In contrast, the cells of the untreated biofilm appeared to be vital and without visible cell damage.

**Figure 6:**
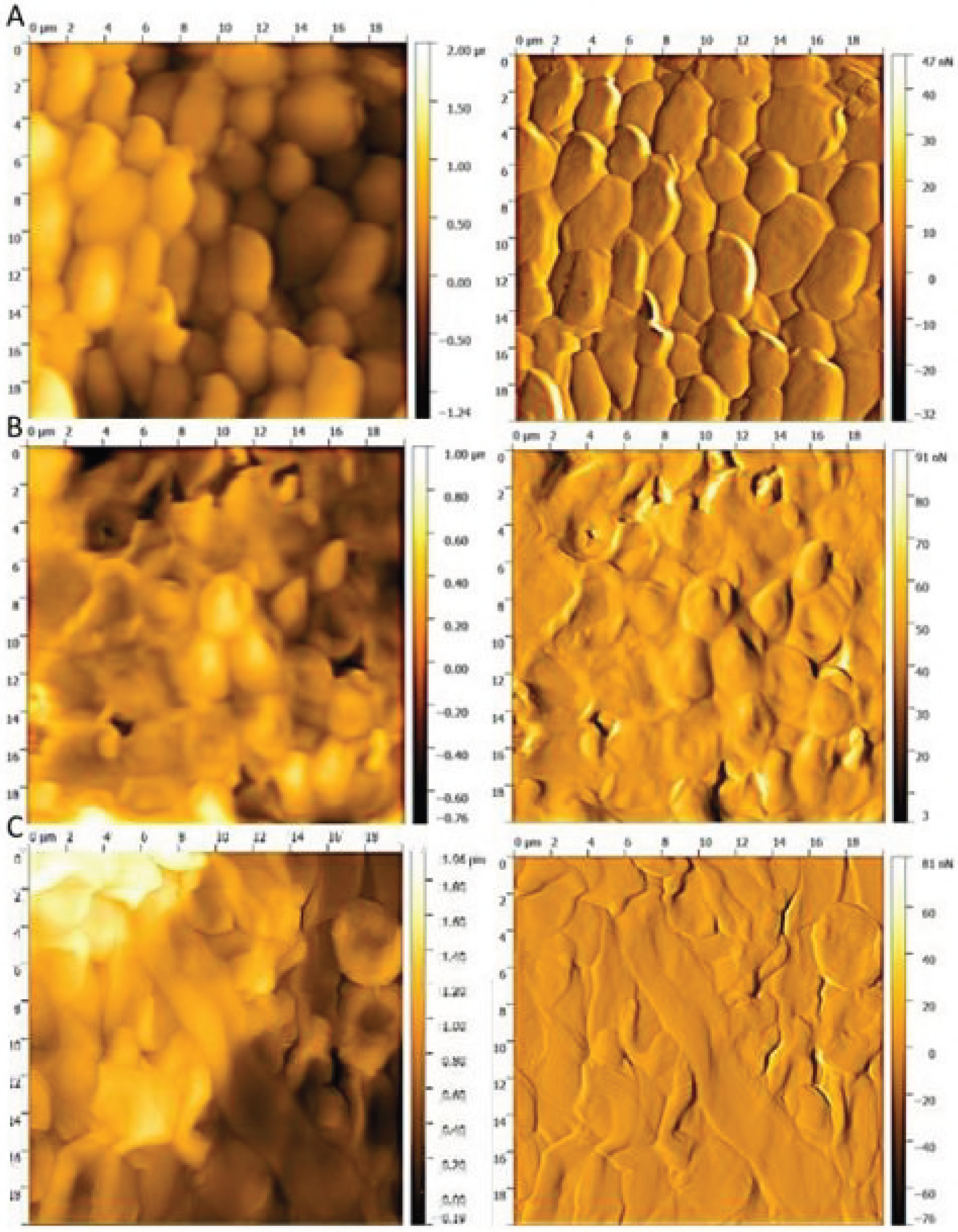
atomic-force-microscopy (AFM) images of the plasma-treated biofilms and references with the kINPen™09, with the topographic image on the left and the error signal image of the same section on the right. A) reference; B) 120 s plasma treatment time; C) 300 s plasma treatment time. The images were taken in contact mode with a cantilever spring constant of k = 0.1 - 0.6 N/m^2^ and a frequency of 0.4 Hz, the set point at 8 N/m^2^ and an area of 20 μm^2^.

## Discussion

Since the invention of non-thermal plasma sources operating at atmospheric pressure, a large number of studies have been published, dealing with the effects of plasma itself (direct mode) and its effluents (indirect mode) on pathogens or living tissue (38–42). However, much less is known about the treatment of living biofilms with non-thermal plasma sources. Koban et al. (43) treated 48 h old *C. albicans* biofilms on titanium discs with different plasma sources and were able to detect an RF of 0.5, after 300 s treatment with the kINPen®09 at a distance of 0.7 mm with 5 slm argon gas-flow. Gorynia et. al. (44) have used the same experimental setup in a distance of 10 mm for 72 h grown *Streptococcus anguinis* biofilms on titanium discs and could also demonstrate a RF of 0.58 after 180 s of plasma treatment. In contrast to these results, we obtained a RF of 1.4 after 60 s of plasma treatment already. Despite the same plasma source has been used, the experimental setup was different. It is therefore difficult to compare our values with the studies previously mentioned. In our study, e.g. no titanium discs were used for cultivation, but *C. albicans* has been grown directly on coated 96-well plates. Furthermore, a distance of 18 mm from the nozzle to the biofilm was selected. Despite the higher distance, we were able to detect a temperature of 54 °C directly in the biofilm after 180 s plasma treatment with the kINPen®09 in the same experimental setup. However, we were able to prove, that the increased temperature has no significante influence on the biofilms. Most likely, the observed difference are due to the use of another *C.* albicans strain compared to the above mentioned similar study. The inactivation of the cells after plasma treatment was proven by different experimental approaches. Counting of CFU`s was used to test for the proliferation ability of treated cells, fluorescence LIVE/DEAD™ assay has been employed to monitor the vitality of the cells and a XTT assay to investigate the plasma effect on cellular metabolism. Using these three complementary methods ensured, that the inactivation of the biofilms by the plasma-treatment was indeed reliable (45).

Recent studies, dealing with the treatment of *C. albicans* biofilms with kINPen®09, demonstrated an inactivation of cells exclusively in the area of plasma effluents (43). However, most of these studies did not investigate the three dimensional inactivation effects within the biofilm. Pei et. al.(46) reported the inactivation of a 25.5 μm thick biofilm of *Enterococcus faecalis* after treatment with a handheld air plasma jet. Here, the authors could demonstrate a complete inactivation of the whole biofilm after 300 s of treatment. Delben et. al.(47) also showed an influence of plasma treatment with an atmospheric pressure plasma jet (APPJ) on three-dimensional biofilms of *C. albicans.* However, both used a different plasma source and a generally different experimental setup. It’s well known, that the biofilm thickness, the microorganism, treatment conditions and the used gas, play a predominant role in microbial inactivation (48).

Even though, a whole series of publications were able to microscopically detect the inactivation of biofilms using plasma treatments, we are not aware of any work that has shown that the bottom of the biofilm was primarily affected by the plasma treatment. This new insights naturally raises the question of how this kind of inactivation was done.

Wilking et. al. (49) were able to detect water channels in *Bacillus subtilis* biofilms for the transport of fluids in biofilms. For this, they used an aqueous solution containing a mixture of fluorescent beads to visualize the connectivity of the channels. Interestingly, they were able to detect a dense network of channels in the center of the biofilm, which reaches to the surface of the biofilm, extends downwards, and spreads over the entire biofilm. Another important aspect of their work was the microscopic proof of the water channel structure. These channels used the surface, to which the cells of the biofilm have adhered, to form their bottom structure. There is a range of other studies that also detect such water channels in bacterial biofilms (50–52). If such water channels exist in Candida albicans biofilms, this could explain the results obtained in our work.

Another explanation for the inhibition of cells, specifically located at the bottom of the treated biofilms could be the rapid diffusion of plasma-relevant reactive oxygen species (ROS) and reactive nitrogen species (RNS) in biofilms. Steward (53) could determining diffusion times of different gases in biofilms. Based on his results a diffusion time of 4.47 s for nitrogen, representing all light gases, could be calculated for the here investigated *C. albicans* biofilms. These results offered plausible explanations for the effects that could be observed in fluorescence microscopy. Another effect that became apparent in fluorescence microscopy was a central spot of living cells, which were visible in the upper layers of 120 s plasma treated biofilms. Desiccation effects, caused by the plasma, could cause this phenomenon. Dead cells on the bottom of the biofilm have disrupted membranes **(Fig. 6)**. As the intracellular fluid escapes, cells might have lost their integrity and detached from the biofilm structure. Due to the dehydration, disruptions and dissolutions in the actin filament may occur (54). As a result, the biofilm structure partially dispersed and the next layer of living cells became visible. This occurs primarily in the center of the biofilm, since the plasma nozzle was placed directly over the center of the biofilm during treatment.

## Conclusion

Microbial biofilms are an important and costly problem, not only in medicine, but also in industrial applications. Thus, an efficient and rapid removal of biofilms is becoming increasingly important. This work provides new insights into the successful inactivation of *C. albicans* biofilms by plasma using the kINPen®09. Notably, this study provides evidences, that the employed plasma treatment caused a primary inactivation of cells located at the basement of the biofilm, which at the same time facilitates the removal of biofilms by other strategies, e.g. mechanical treatment. This novel insight into plasma inactivation of biofilms indicates that plasma is well-suited to fight biofilms growing on various surfaces, since surface detachment of biofilms is an important prerequisite for an effective removal of biofilms, especially in industrial settings.

## Materials and Methods

**Fungal strain and growth conditions.** A fungus strain with primarily vertical biofilm growth named *Candida albicans* SC5314 (ATCC® MYA-2876™) was used. The fungus was grown on Sabouraud agar and incubated 24 h at 37 °C. A suspension of the fungus in PBS (pH 7.2 according to Sörensen) was adjusted to an OD of 0.375 at 520 nm. 1 ml of this suspension was added to 9 ml RPMI medium (without bicarbonate, Merck, Darmstadt, Germany), which served as an inoculum for biofilm cultivation in microtiter plates. For plasma treatment, a coated Polysterol 96-well plate (Sarstedt, Nümbrecht, Germany) was used. As an exception, a 12-well plate was used for the AFM. 200 μl of the suspension were pipetted per well and incubated 90 min at 37 °C and 80 rpm on a rotary shaker to allow cell attachment. After incubation, the medium was removed and the well washed with 200 μl PBS (pH 7.2 according to Sörensen) and 200 μl RPMI medium was added. The plate was incubated 24 h at 37 °C and 80 rpm on a rotary shaker, resulting in well-established *C. albicans* biofilms for further analysis. This protocol was kindly provided by the research group of Dr. Christiane Yumi Koga-Ito, Institute of Science and Technology ‒ UNESP, Department of Oral Sciences and Diagnosis.

**Plasma source.** kINPen®09 (neoplas GmbH, Greifswald, Germany), a commercial APPJ, was used for plasma treatment. Using a DC power supply (system energy: 8 W at 220 V, 50/60 Hz) and 99.99 % argon gas connection (Linde, Pullach, Germany) with 5 slm, a plasma was generated under atmospheric pressure. The inside of the APPJ handpiece consists of a 1.6 mm diameter quartz capillary with an embedded electrode of 1 mm diameter **(Fig.7)**. When the APPJ was in operation, a high-frequency voltage of 1.1 MHz, 2-6 kVpp was applied to the electrode. This generated the plasma at the tip of the electrode and expelled it into the surrounding medium forming an effluent (55).

**Figure 7:**
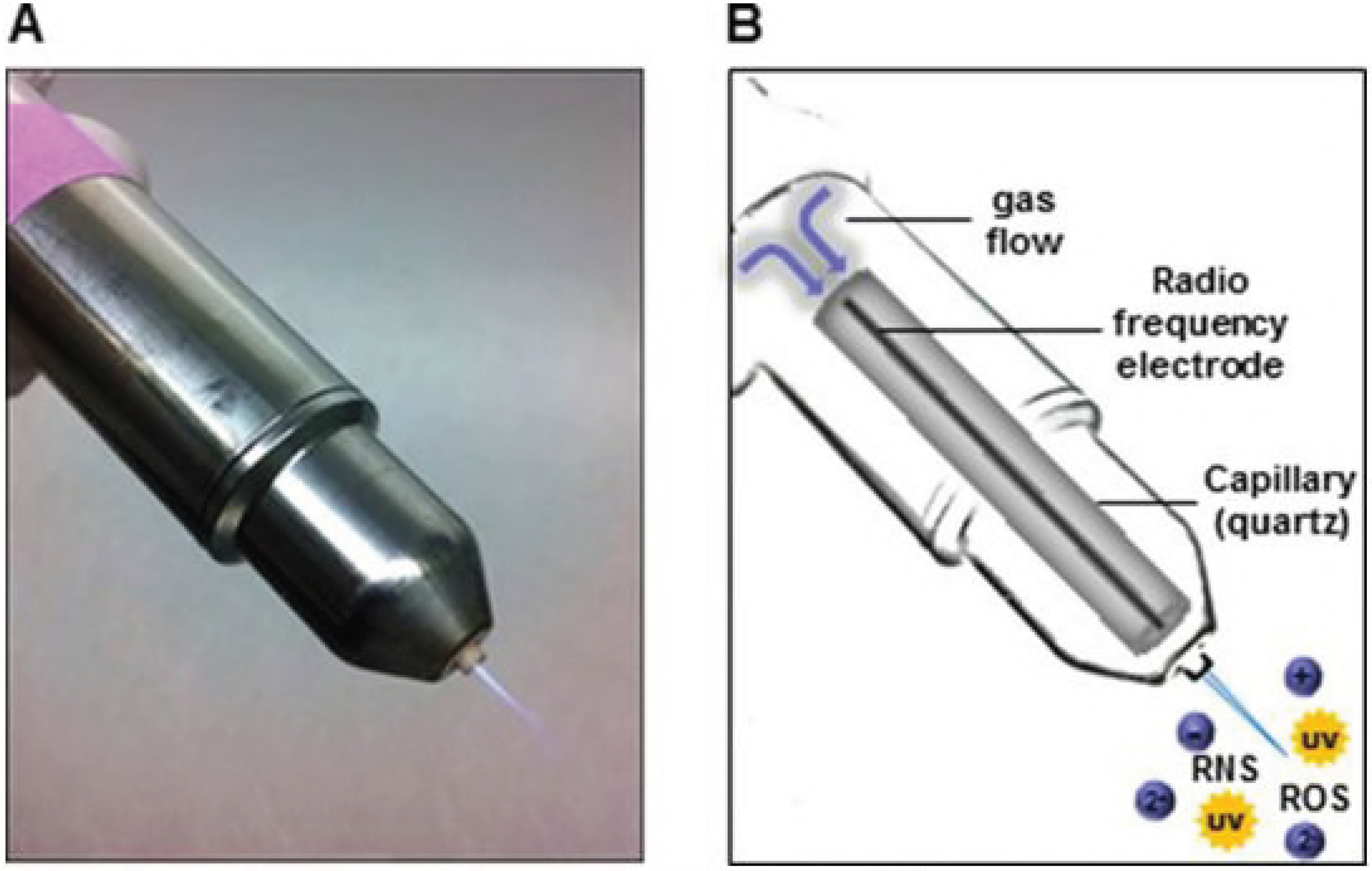
A) Plasmajet (APPJ) kINPen™09 (Neoplas GmbH, Greifswald, Germany); B) schematic structure of the APPJ and composition of the physical plasma (Weiss,2015).

**Plasma treatment of *C. albicans* SC5314 biofilms.** Established biofilms were washed with PBS (pH 7.2 according to Sörensen), followed by complete removal of the liquid. Five different time intervals were used at a distance of 18 mm from the plasma effluent tip to the biofilm surface using a computer-controlled XYZ table (CNC Maschinenbau, Geldern, Germany), to which the plasma source was attached. Only one sample group was treated per time to avoid dehydration effects of the biofilms. For CFU counting, fluorescence LIVE/DEAD™ assays and XTT assays, 200 μl PBS (pH 7.2 according to Sörensen) were added per well, the biofilms were mechanically disrupted and the resulting cell suspension was collected. This step was repeated three times, until the complete biofilm was transferred.

**Analytical determinations.** First, the effect of the gas flow on the biofilms was investigated and the temperature of the biofilms was measured directly. 5 slm argon gas was used for the gas flow measurements as it was the case with the following plasma treatments. However, the gas was not ignited. For temperature measurements, biofilms were plasma treated and thermal images were taken directly after the treatment with a FLIR thermal imaging camera (FLIR Systems, Frankfurt am Main, Germany) in a distance of 20 cm. Temperature and gas flow on their own have no significant influence on biofilms (data not shown). CFU assay was used to determine the proliferation rate of the *C. albicans* biofilms after plasma treatment. Serial dilution of the samples was done by diluting the sample solution after plasma treatment 1:10 with VF-water. The references were finally diluted 1:10,000 and the samples 1:1,000. Each dilution step was plated on Sabouraud agar by pipetting 10 μl per dilution onto the plate and by means of tilting technique, the sample was spread out. The plates were then incubated for 24 h at 37 °C. The colonies for the respective dilution levels were counted and the CFU×ml^−1^ were calculated representing 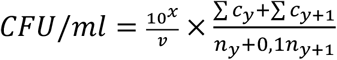 where 10^x^ is the dilution factor for the lowest dilution, v is the volume of diluted cell suspension per plate in ml, Σc_y_ is the total number of colonies on all (n_y_+1) plates of the lowest evaluated dilution level 10^−x^ and Σc_y_+_1_ is the total number of colonies on all (n_y_+1) plates of the next highest dilution level evaluated 10^−(x^+^1)^. After calculating the CFU×ml^−1^, the reduction factor (RF) could be determined using RF = MV_r_log_10_ − MV_s_log_10_, where MV_r_log_10_ representing the mean value of the CFU×ml^−1^ of the reference group and MV_s_log_10_ is the mean value of the CFU×ml^−1^ of the treated samples. The confidence range (CR), which was used for the calculation of the diagram error bars, was determined using the formula 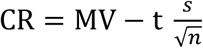, where t is the t value from the t distribution table, s is the standard derivation and n is the number of replicates. The experiments were repeated fourfold with n = 6.

**Fluorescence LIVE/DEAD™ assay.** The LIVE/DEAD™ BacLight™ Bacterial Viability Kit was prepared by mixing reagent A and B at a ratio of 1:1. After the mixing step, 0.9 μl of this solution was pipetted to 300 μl of the sample solution after plasma treatment for each well followed by incubation for 20 min at room temperature with 80 rpm on a rotary shaker in the dark. In the next step, the 96-well plate was scanned with an excitation wavelength of 470 nm and emission wavelength of 530 nm for green fluorescence and 630 nm for red fluorescence with the Varioskan-Flash® device (Thermo Scientific, Waltham, USA). At least, ratio G/R was calculated by dividing the adsorption value of red fluorescence from the value of green fluorescence. The experiments were repeated fourfold with n = 6.

**XTT assay.** For the XTT assays, XTT Cell Proliferation Assay Kit (Applichem, St. Louis, USA) was used. The activation solution and the XTT solution were mixed 1:50 and 50 μl were transferred in a 96-well plate containing 100 μl sample solution after plasma treatment per well. The 96-well plate was incubated for 2 h at 37 °C, 80 rpm in the dark. After the incubation time, 96-well plate were scanned at a wavelength of 470 nm using a Varioskan-Flash® device. The obtained values were blank corrected. The experiments were repeated fourfold with n=6.

**Fluorescence microscopy.** Biofilms were cultivated as previously described, using a black 96-well plate with a glass bottom (PerkinElmer, Hamburg, Germany). Plasma treated biofilms were resuspended in 300 μl of 0.85 % NaCl after treatment to avoid dehydration of the biofilms. The LIVE/DEAD™ BacLight™ Bacterial Viability Kit was prepared by mixing reagent A and B in a ratio of 1:1. Next, 0.9 μl of the fluorescence solution was added to each well and the 96-well plate was incubated at room temperature for 20 min and 80 rpm in the dark. After incubation, the supernatants were displaced and the samples were washed again three times with 0.85 % NaCl. Epifluorescence images were acquired using Operetta CLS High-Content Imager (PerkinElmer, Hamburg, Germany) using the following objectives (Zeiss, Oberkochen, Germany): 1.25x (air, numerical aperture (NA) = 0.03), 5x (air, NA = 0.16), 20x (air, NA = 0.4), 40x (washing, NA= 1.1). Depending on the experiment, several fields of view were recorded and combined in the software. SYTO™9 was excited by 475 nm (110 mW) LED and the fluorescence was collected with a 525 ± 25 nm band pass filter. Propidium iodide was excited by 550 nm (170 mW) LED and fluorescence was collected with a 610 ± 40 nm band pass filter. A laser autofocus (785 nm) was available for all measurements. The images were displayed using Harmony 4.6 software.

**CLSM.** Biofilms were cultivated, plasma treated and LIVE/DEAD™ stained as described above. After the staining and washing procedure, the supernatants were removed and the biofilms were analyzed using a Zeiss LSM 510 microscope (Carl Zeiss, Jena, Germany) equipped with a 10x objective (air, NA =0.1). Filter and detector settings for monitoring SYTO™9 and propidium iodide fluorescence (excitation at 488 nm using an argon laser, emission light of SYTO™9 selected with a 505 - 530 nm bandpass filter, emission light of propidium iodide selected with a 650 nm longpass filter). Three-dimensional images were acquired using the ZEN 2009 software (Carl Zeiss, Jena, Germany) with an area of 1272.2 μm × 1272.2 μm and z-stack sections of 5.5 μm.

**Atomic-force microscopy.** For better adhesion of the coverslips (13 mm, Sarstedt, Nümbrecht, Germany) to the surface of the well plates and to avoid growth of the fungus at the bottom of the coverslips, 50 ml Gelrite™ (Duchefa, Haarlem, Netherlands) was autoclaved and directly used after the autoclaving process due to a rapid thermal curing process. A volume of 200 μl liquid Gelrite™ were pipetted to each well of a 12-well plate. The coverslips were placed at the surface of the liquid Gelrite™ and were thermally cured. The biofilms were cultivated as described above and 1 ml of the RPMI medium were pipetted to each well until the coverslips were topped with the medium. 12-well plates were incubated at 37 °C for 1 h and 80 rpm. After the incubation time, additional washing steps with 1 ml of 0.85 % NaCl each well, were followed. On 24 h later, the cultivation medium was removed and the biofilm overgrown coverslips were washed with 1 ml 0.85 % NaCl again. After washing, samples were treated for 120 s and 300 s by the kINPen®09 plasma effluent. Dehydration of the coverslips before AFM analysis was avoided by using a humidity chamber. The AFM measurements were carried out on a DI CP II SPM (Veeco, Plainview, USA), which was mounted on a vibration-free object table (TS-150, TableStable, Zwillikon, Switzerland). The setup was standing on an optical bench encased by an additional acoustic protection. The AFM was equipped with a linearized piezo scanner, on which the coverslips were mounted on a metal sample holder with leading tabs. The samples were measured using cantilevers with nominal spring constants of k = 0.1-0.6 N×m^−2^ in contact mode. The pictures were taken by a scanning speed of 0.4 Hz by a picture size of 20 μm^2^ and a set point = 8 N×m^−2^. Pictures were edited with Gwyddion (Czech Metrology Institute, Brno, Czech Republic).

## Acknowledgment

This work was supported by the Leibniz Institute for Plasma Science and Technology, INP in Greifswald, Germany, and the DFG-TRR34 (A3 subproject granted to Katharina Riedel). Fluorescence microscopy was supported by the group of Sander Bekeschus within the project “Zentrum für Innovationskompetenz plasmatis, Nachwuchsgruppe: Plasma-Redox-Effekte”, which was financially supported by the German Federal Ministry of Education and Research (BMBF).

